# The Role of Signal in Producing Saltatory Approach to Cryptic Stationary Targets

**DOI:** 10.1101/2025.06.30.662452

**Authors:** Eden Forbes, Peter M. Todd, Randall Beer

**Affiliations:** Cognitive Science Program, Indiana University, Bloomington, Indiana, USA; Department of Psychological and Brain Sciences, Indiana University Bloomington, Indiana, USA; Neuroscience Program, Indiana University, Bloomington, Indiana, USA; Department of Informatics, Indiana University, Bloomington, Indiana, USA

**Keywords:** foraging, pursuit, animal movement, saltatory movement, agent-based model

## Abstract

Saltatory movement strategies in animal search are well known, but their role in pursuit and localization of targets is less so. There are multiple features of forager-target interactions that may generate saltatory forager movement during pursuit and/or localization of the target. Here, we explore how the reliability of the target’s signal and the dependence of the forager’s perception of that signal on the forager’s own motion can generate a variety of different continuous and saltatory pursuit patterns in evolved model foragers with stationary targets. We use dynamical analyses of evolved forager nervous systems to show the possible mechanisms through which saltatory movement was generated and how signals influence those movements in a foraging animal. Saltatory movement during pursuit independent of signal was routinely found when the forager’s perception depended on its movement. Saltatory movement patterns could also be generated in otherwise cruise-like behaviors when target signal either excited or inhibited ongoing forager movement. Additionally, the magnitude of the signal’s influence was found to depend on proximity to the target during pursuit. These models present hypotheses for future empirical research and emphasize the importance of exploring variation in animal movement during different phases of foraging.

## Introduction

Increased fidelity and quantity of animal movement data have facilitated a more coherent field of animal movement and the ecological circumstances that shape it (Nathan et al., 2008).

Often, animal movement is examined in the context of foraging problems, for instance asking what movement patterns yield the greatest or most energy efficient encounter rates with targets in the environment (MacArthur & Pianka, 1966; Pyke et al. 1977; Gerritsen, 1984). Early work in this vein identified a clear dichotomy of foraging mode between animals that move continuously while foraging (“cruise”) and those that wait to attack passing prey (“ambush”) (Pianka, 1966; Huey & Pianka, 1981; McLaughlin, 1989). However, animal movement data rarely falls cleanly into unimodal or bimodal distributions around these two foraging modes (Cooper, 2005). Cruise and ambush patterns are extremes on a spectrum of movement strategies. In between lies a range of intermediate stop-and-go or saltatory movement patterns (Evans & O’Brien, 1988; O’Brien et al. 1989) that are ubiquitous across taxa (O’Brien et al. 1990). Saltatory movement encompasses a wider variety of behavior than the simple cruise/ambush dichotomy, as well as variations in that movement according to prey size (O’Brien et al. 1989), habitat complexity (Brownsmith, 1977; Ehlinger & Wilson, 1988), and prey density (Moermond, 1979).

That said, to this point most emphasis has been put on large-scale search processes while glossing over other parts of the foraging cycle (Holling, 1966; Lima & Dill, 1990) including pursuit, capture, and consumption. Many animals certainly spend most of their time searching, whether for a single target or a preferred target amongst many. As such, animal movement data is often best suited to characterize search, as it yields the largest amount of data. But focus on search alone overlooks the problem of signal reliability post-encounter; an encounter with a target signal does not imply high fidelity or continuous information during pursuit of that target. Models that focus solely on the cruise/ambush dichotomy (e.g., Scharf et al. 2006; Higginson & Ruxton, 2015) or those few that include saltatory strategies (e.g., Benichou et al. 2011; Gurarie & Ovaskainen, 2013) thus do not explore movement strategies post-encounter.

This is a particular problem given that most targets of foraging animals do not want to be found. Most prey exhibit a form of behavioral crypsis in the presence of predators. Examples of behaviorally cryptic prey abound even when that prey is sessile and has no means of outright escape. For example, embryos of cuttlefish (Bedore et al. 2015), ray (Ball et al. 2016), shark (Kempster et al. 2013), and skate (Gervais et al. 2021) cease active oxygen filtration of their egg capsules (Long & Koob, 1997) or other activity that creates detectable chemical traces in the presence of predators (Du et al. 2022). Embryonic shorebird acoustic calls similarly decline in the presence of predator calls (Kostoglou et al. 2021). So, even in seemingly simple pursuit situations, foraging animals often face the challenge of unreliable signal from their targets. This challenge is relevant to all perceptual modalities (Ruxton, 2009) and applies to almost all animal taxa (Stevens & Ruxton, 2019).

Perception of targets is a known determinant of foraging mode during search (Rice, 1983; de Queiroz, 2003; Torres et al., 2017; Stöckl & Kelber, 2019; López-Cruz et al., 2019; Kelkar et al., 2018) but is often assumed to be constant and continuous in both search (e.g., Koopman, 1956; Gerritsen & Strickler, 1977) and pursuit (e.g., Nahin, 2012; Weintraub et al. 2020) especially in models. Cryptic targets violate that assumption. Furthermore, perception and movement on the part of the forager often involve direct trade-offs. Some behaviors necessary for perception require slowing or cessation of movement. Mechanoreception, olfaction, audition, and electroreception all require active reception of stimuli through behaviors such as whisking (Hartmann, 2001), sniffing (Wachowiak, 2011), and electrolocation (Nelson & MacIver, 2006; Caputi, 2023) that can be hindered by motion. Motion can also decrease perceptual sensitivity due to introduced noise, evident in suppression of self-generated signals by animal nervous systems (Blakemore et al., 1998; Foo & Mason, 2005). These trade-offs between perception and movement are likely amplified in circumstances with cryptic targets.

We know very little about the mechanisms relating signal perception to movement patterns in foraging animals as mediated both by how animals receive signals and the reliability of those signals. Here, we present a model system that demonstrates how reliability of target signal and perceptual modality each and in combination elicit a variety of continuous and saltatory movement patterns during pursuit in a physiologically standardized model forager. We outline how foragers were evolved (see below) to solve foraging problems with reliable or unreliable signals and with or without perceptual dependence on movement. We then describe the various mechanisms by which unreliable signal or movement-dependence yielded saltatory patterns of motion in our model foragers. After measuring the magnitude and sign of various behavioral responses to signal, we summarize a variety of strategies that emerged in our evolved populations. Finally, we compare those strategies to some of the existing behavioral data and concepts in the literature to demonstrate their applicability. We hope this contribution will help researchers better diagnose the signals being used by their study species in pursuit problems and will progress our understanding of animal movement more broadly.

## Methods

### Task Environment & Forager Agents

Model agents (henceforth, foragers) are situated in a wrap-around 2D arena which can have any side length greater than their detection radius *d* (Figure 1A). Foragers and their targets are approximated by points in that arena with Euclidean coordinates that change depending on forager movement during simulation. Foragers move depending on the output from a model nervous system described as a continuous-time recurrent neural network (CTRNN, Figure 1B). These networks, which can be sensitive to both environmental circumstance and internal state, have been used in many studies of adaptive behavior given their approximation of real neural dynamics (Beer & Gallagher, 1992; Beer, 1995a; Beer 1995b). Change of each CTRNN node’s state over time in a network with *N* nodes and *S* sensors is given by the equation:

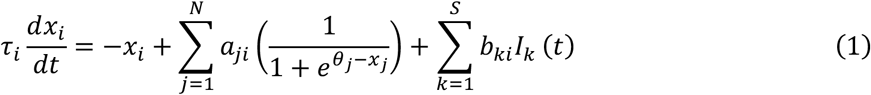

**Figure 1:**
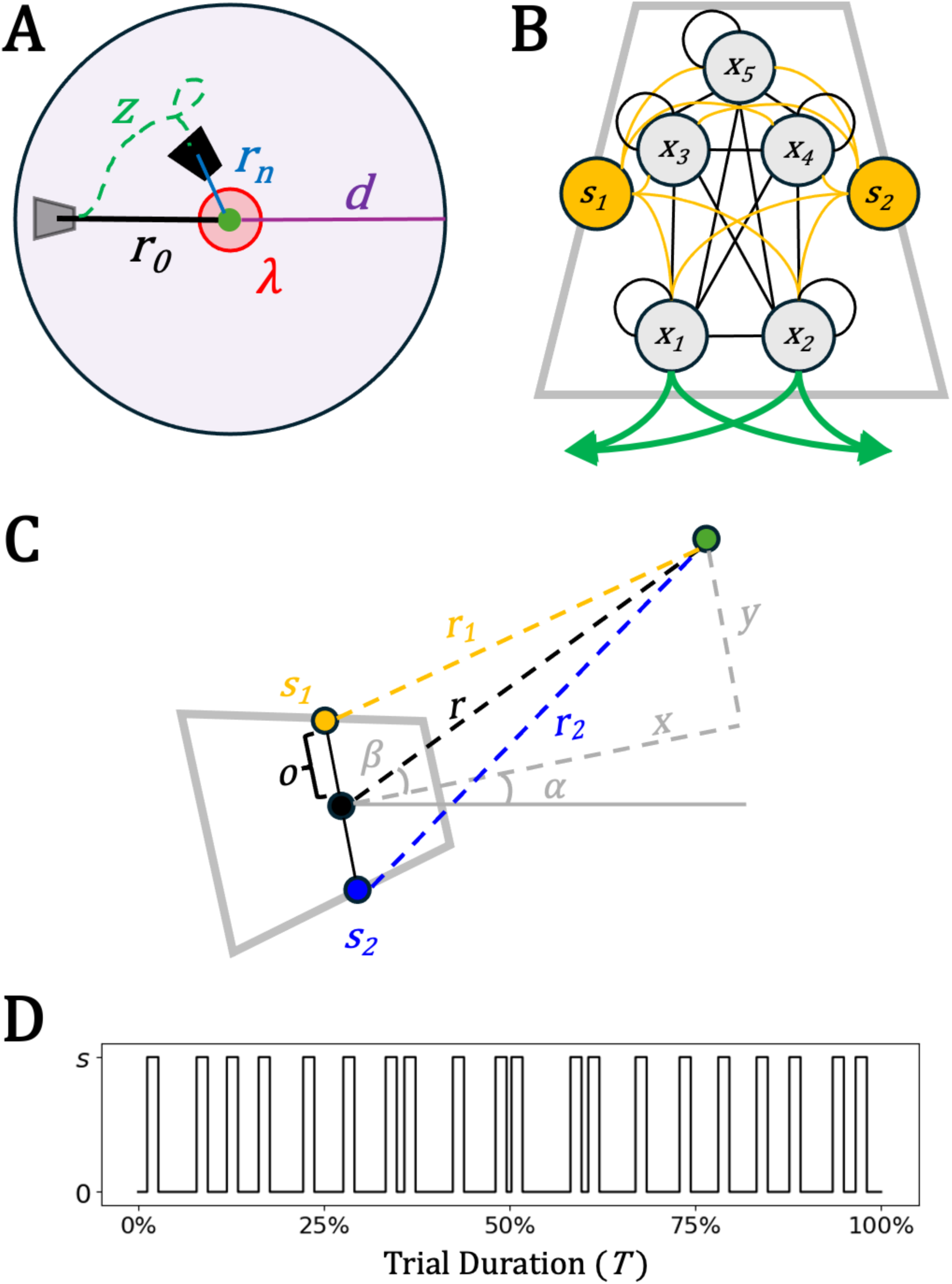
Task environment, nervous system, and sensor arrangement of forager agents. **(A)** After a transient period, a target (blue dot) is put within detection radius d of the agent (grey rhombus) at initial distance r0. Trial fitness is calculated depending on the forager’s (black rhombus) final distance from the target rn, with additional bonuses for ‘attacking’ (being within radius λ). **(B)** Pictural representation of the fully connected 5-node CTRNN and the two fully connected sensors. Connections within and between nodes and sensors may be either excitatory or inhibitory depending on the outcome of the evolutionary algorithm. Motor output is determined only by the states of x1 and x2. **(C)** Sensors are offset (o) from the center of the agent. As such, the distances relevant to the two sensors (r1 and r2) are calculated according to the current distance r, the current heading *α* and the current angle to target *β* (eq. 4). **(D)** Sample stimulus patterns from an unreliable target across an entire trial with sI = 20 and sP = 0.3.

where *x_i_* is the node’s current state, 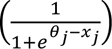 is a sigmoidal shaped firing frequency curve governed by the firing threshold or bias *θ*, *τ* is a time constant that determines the rate of state change, *α_ji_* is the strength of the connection from the *jth* to the *ith* node, and *I_k_* represents the external input from the *kth* sensor scaled by sensor weights *b_ki_*. Movement of the forager in terms of velocity (*v*) and torque (*q*) is determined by relations between outputs of nodes *x_1_* and *x_2_* of the CTRNN:

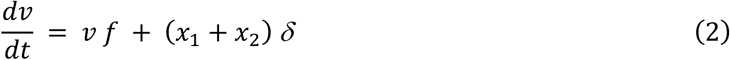

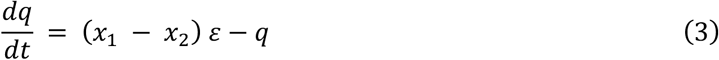

where *f* is a friction coefficient presumed to be less than 1, δ is the maximum possible thrust, and *ε* is the maximum possible turning angle. Maximum *v* (given by setting *x_1_* and *x_2_* = 1; eq. 2) is *2δ/f*.

Foragers have two sensors displaced by offset *o* on the left and right side of its body that contribute to nervous system inputs *I*. These sensors’ values *s_1_* and *s_2_* are determined by distance from the target, according to the geometric translation:

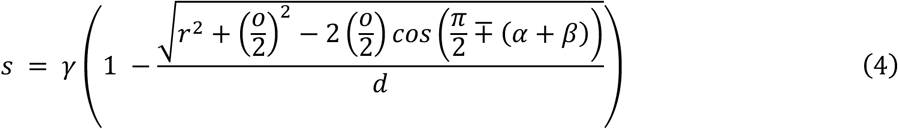

where *γ* is a scalar that determines maximum sensor value, *r* is the current distance from the forager to the target, *o* is the offset of each sensor from the agent’s center, *α* is the agent’s current heading (relative to 0 degrees) and *β* is the angle from the agent’s heading to the target (Figure 1C, D). Whether or not the heading and relative angle are added or subtracted in the cosine function delineates the left-side and right-side sensors (*s_1_* and *s_2_*).

Each of the equations that define the perception and movement of foragers (eq. 1 – 4) are calculated each twentieth of a timestep for the total simulation duration (*T* timesteps). So, the stepsize between each calculation, or the *dt* of each equation here, is 0.05. That fine resolution allows an otherwise discrete-time simulation to be analyzed using continuous methods for differential equations (see below).

### Signal Availability and Trial Structure

Two signal features were manipulated to create four pursuit problems. First, signal from the target was either reliable (*R*) or unreliable (*U*). Reliable signal means that the target was perpetually creating stimuli available to the forager. Unreliable signal implies that the target would start and stop some behavior that is creating those stimuli. Here, unreliable targets provide signal for a random percentage *s_P_* of each of some number of intervals *s_I_* during a trial (e.g., Fig. 1D). Resulting signal lengths from unreliable targets are thus determined by *s_P_ * T/ s_I_*. Second, the ability of the forager to detect signal was either independent of movement (*I*) or dependent on movement (*D*). To impose movement dependence, *D* foragers were unable to detect any signal if their velocity *v* rose above a threshold *Ø*. In combination, signal reliability and movement dependence created the four pursuit problems: reliable movement-independent signal *RI*, unreliable movement-independent signal *UI*, reliable movement-dependent signal *RD*, and unreliable movement-dependent signal *UD*.

Foragers in each condition were tested on 24 trials per generation of evolution (see below for evolutionary algorithm). At the start of each trial, the CTRNN states are randomized (0 – 1). After a short transient period (*T_T_* timesteps), an ‘encounter’ occurs and a target is placed somewhere inside the forager’s detection radius *d* at an initial distance *r_0_* randomly between 3 – 7 units less than *d* (here, *d* = 29). The 24 encounters were initialized with the target on each π/24 interval between the direct left and direct right of the agent (targets were assumed to never appear behind the forager). Trials ran to maximum duration *T* and were ended before *T* only if the forager was within an interaction distance of the target (*r_n_* ≤ λ) and had come to a stop (*v* ≤ *Ø*) thus ‘attacking’ the target. That stopping condition further necessitates behavior that is oriented to the target, instead of those that may focus on randomly searching a tight area. Performance during a trial was determined by a simplified version of a chemotaxis index (e.g., Izquierdo & Lockery, 2010) which measures the relative change in distance between forager and target at the start and end of a trial. Successful pursuit, those trials that ended close enough to the target to attack (*r_n_* ≤ λ), were given additional bonuses according to the efficiency of their route length *z* and how quickly they accomplished the trial. Failed pursuit, trials that ended outside of detection radius (*d* ≤ *r_n_*) and had thus lost the target, were given a fitness of 0. In sum, for each trial, forager fitness *F_F_* was evaluated according to:

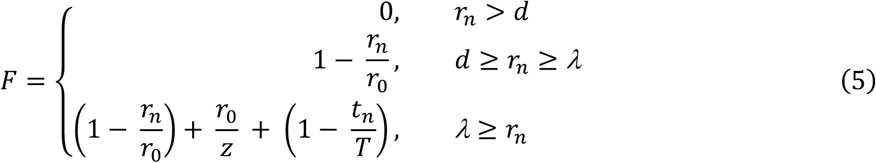

where *r_n_*. is the distance between the forager and the target at the end of the trial, *z* is the total distance traveled by the forager, and *t_n_*. is the time taken to complete the trial. Final fitness for a generation of the evolutionary algorithm was determined by calculating the mean across all 24 trials 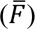. Specific parameters for all evolutionary algorithms, task environments, and CTRNN value distributions in evolution can be found in the Appendix (Section 1).

### Evolutionary Algorithm

A real-valued genetic algorithm (Mitchell, 1998) was implemented for 100 populations of foragers in the four pursuit conditions to generate sets of behavioral solutions for each pursuit problem. The genetic algorithm had access to CTRNN biases (*θ*), time constants (*τ*), connection weights (*α*_ji_), and sensor weights (*b*_ki_). Our genetic algorithm was generational with rank-based selection. During evolution, any gene may take on values from [-1, 1] which are then deterministically translated to phenotypic parameters. That said, the initial population of each evolutionary run was seeded with genomes with random gene values from [-0.01, 0.01] except for *θ_1_* and *θ_2_* which were set to −1. This constraint was imposed to aid the evolution of movement-dependent (*D*) foragers that need to stop moving to detect signal. Because foragers with low motor neuron biases are generally stationary, a more stationary initial population helps movement-dependent populations converge on viable intermittent movement solutions rather than exploring a massive amount of parameter space that will never detect any signal (where *v* is always greater than *ø*). Seeding populations this way had no obvious effect on the final genetic distributions of movement independent (*I*) foragers. Successive generations were created by applying random Gaussian mutations to high performing genomes and a small chance of uniform crossover between those same genomes. A small fraction of those best performers from the previous generation were left unchanged.

## Results

### Evolution, Robustness, and Movement of Foragers in Pursuit

Successful attack of the target (ending the trial with *r_n_* ≤ λ), will always yield a fitness close to 1 with some additional bonuses for route efficiency and time efficiency. As such, we determined successful foragers to be those that exceeded mean fitness of 1 across all trials (F > 1). Successful pursuit strategies were found for all four combinations of signal reliability and movement dependence. The best forager from 86/100 *RI* runs, 46/100 *UI* runs, 69/100 *RD* runs, and 37/100 *UD* runs met that threshold. Best fitness was consistently lower when target signal was unreliable (*UI* and *UD* foragers) than when it was reliable (*RI* and *RD* foragers; Table 1). This is unsurprising; foragers with less information available to them face a more difficult task. That said, movement dependence (*UD* and *RD* foragers) did not appear to reduce best possible fitness, although there was more variance in the outcome of evolutionary runs. We conducted a robustness analysis on the best performing agent from each pursuit task, evaluating their performance at thousands of combinations of initial position and initial heading to further verify the F > 1 criterion (Appendix Section 2). Notably, these best strategies were not guaranteed to succeed from every initial condition in the trained area. While foragers with reliable (*R*) signal were almost universally successful, even the best foragers with unreliable (*U*) signal availability regularly failed. Still, real animals are not entirely successful in pursuit either. Having generally successful foragers in all four conditions (*RI, RD, UI, UD*) allows us to directly compare the movement patterns that are shaped by those pursuit problems.

**Table 1:** Statistical summary of evolutionary performance and resulting strategies in each pursuit condition. Note here that percentages are nested. For example, 98.6% of RD foragers are periodic samplers. The subsequent breakdown into signal chasing, listening, and mixed strategies is based on that subgroup, thus adding up to 100%.

| Strategy | RI | UI | RD | UD |
| --- | --- | --- | --- | --- |
| EA success | 86% | 46% | 69% | 37% |
| Best final EA fitness | 2.106 | 1.440 | 1.593 | 1.215 |
| Mean final EA fitness ($\bar{F} > 1$) | 1.669 | 1.094 | 2.076 | 1.685 |
| % Periodic sampling ($\bar{F} > 1$) | N/A | 0% | 98.6% | 100% |
| % Signal chasing | N/A | - | 56.5% | 45.9% |
| % Signal listening | N/A | - | 11.6% | 10.8% |
| % Mixed strategy | N/A | - | 31.9% | 43.5% |
| % Non-periodic sampling ($\bar{F} > 1$) | N/A | 100% | 1.4% | 0% |
| % Signal chasing | N/A | 2.2% | 100% | - |
| % Signal listening | N/A | 67.3% | - | - |
| % Mixed strategy | N/A | 30.5% | - | - |

Solutions for all four pursuit problems yield unique behavioral features despite evolving to solve a fundamentally similar task. Qualitatively, we examined the movement patterns of a subset of the best evolved foragers by measuring their movement with a target kept at a constant distance. Each task yielded different movement patterns according to the cruise-saltatory-ambush schema introduced by O’Brien et al. (1990) (Figure 2). *RI* foragers, those that match the common assumption of continuous perception during pursuit, almost universally demonstrated cruise behaviors with close to constant velocities. Conversely, both movement-dependent conditions (*RD* and *UD*) demonstrate saltatory patterns of movement with various frequencies and durations of stationary and mobile periods. *UD* foragers evolved solutions both with and without regular periods in their movement, while *RD* foragers were all regularly periodic. *UI* foragers demonstrated a broad range of movement solutions, with the best being saltatory but without a regular period.

**Figure 2:**
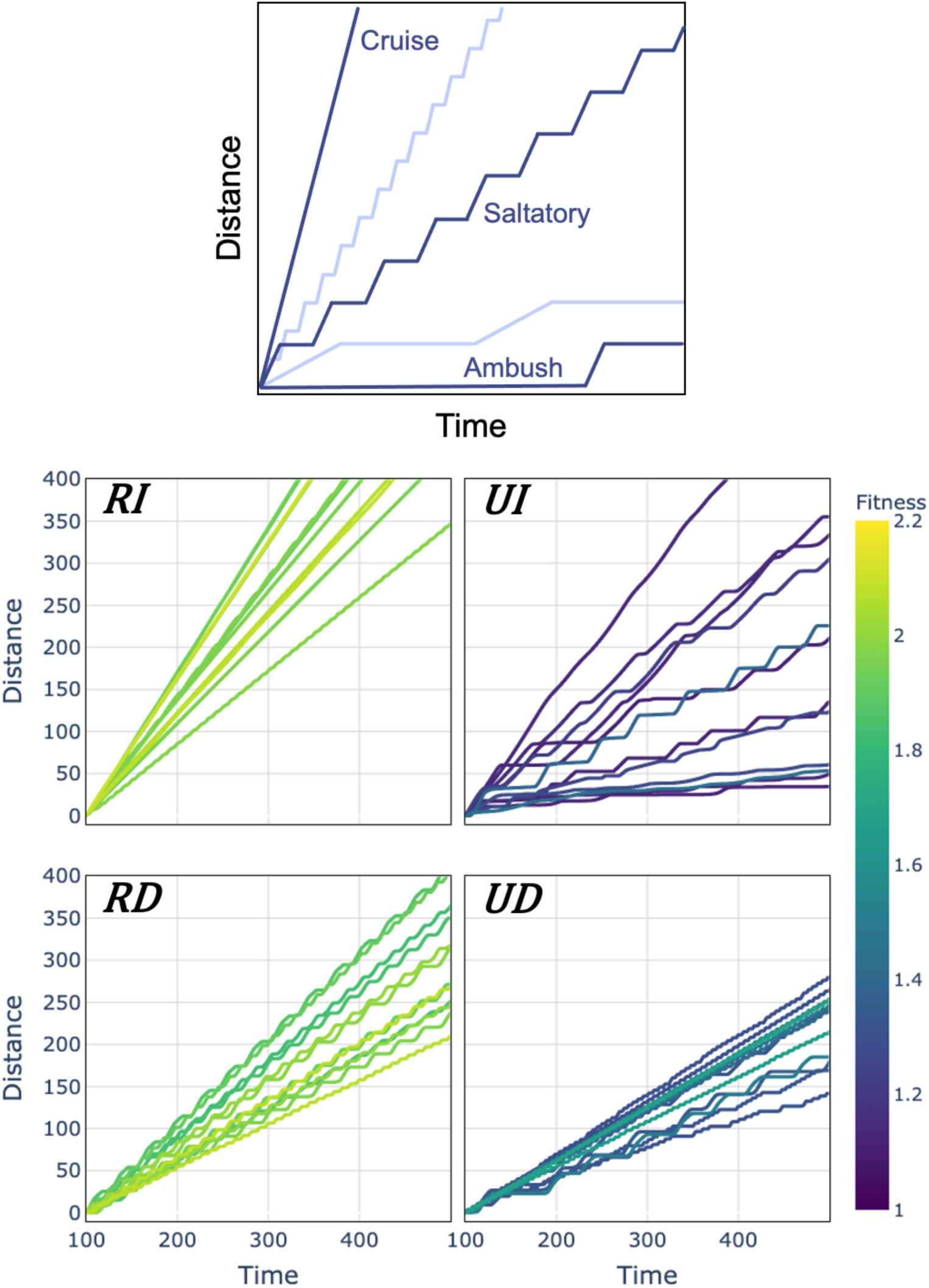
Movement behaviors from the top 12 foragers evolved for each of the four pursuit tasks. The top panel shows archetypal animal search modalities, adapted from O’Brien et al. (1990). Cruising foragers move constantly while their ambush counterparts only move to strike passing prey. However, most organisms’ movement falls somewhere in between and is characterized by stop and go or saltatory patterns. A variety of modalities were also found in the pursuit task. Foragers with reliable signal and movement independent perception **(RI)** demonstrated cruise-like behavior with little variation in velocity. Foragers with unreliable signal and movement independent perception **(UI)** exhibited a wide range of pursuit strategies, including many saltatory strategies. Foragers with reliable signal and movement dependent perception **(RD)** relied entirely on saltatory behavior to successfully pursue targets. Foragers with unreliable signal and movement dependent perception **(UD)** similarly relied on saltatory strategies but with two different apparent kinds, those with frequent short stops and those with long samples and stops.

### Dynamical Mechanisms of Saltatory Movement with Signal

While *RD*, *UI*, and *UD* foragers all exhibited saltatory patterns of motion, the mechanisms underlying that saltatory motion and variation in that motion differed. We identified three mechanisms that drive saltatory motion in our evolved populations, each of which are differently caused and/or shaped by target signal during the task: periodic sampling, signal listening, and signal chasing. These mechanisms were identified by examining the behavior of all best performing foragers with F > 1 in the *UI*, *RD*, and *RI* conditions.

#### Periodic Sampling

Evolved foragers either moved with or without a regular period. Foragers with a regular cycle of acceleration and deceleration exhibited saltatory patterns of movement independent of target signal, but the duration and in turn distance of that movement could be shaped by the received signal. Identifying what differentiates periodic versus non-periodic samplers requires examining the equilibria of the nervous system dynamics of token foragers. Because CTRNNs are described by ordinary differential equations, their behavior is amenable to dynamical systems analysis. The dynamics of periodic sampler nervous systems are cyclical without any signal. That is, a stable limit cycle in the nervous system dictates movement that changes from fast (upper right points) to slow (lower left) and back in the absence of input (Figure 3A, top left). Periodic samplers only evolved in cases with movement-dependent perception (*RD* and *UD* foragers), given that those foragers necessarily had to balance a trade-off of signal only being available at low velocities but also needing to move considerable distances to pursue targets. An intrinsic limit cycle allows periodic perception of the environment by the forager, as velocity regularly dips below the perceptual threshold (*v* ≤ *Ø*). If signal is never introduced this pattern occurs at a fixed period indefinitely. However, when a sample overlaps with available signal, new attractors at that signal level shift the nervous system’s attractors away from that baseline cycle, pulling nervous dynamics off the signal-absent limit cycle (Figure 3A, middle left). The location of those attractors relative to the original cycle determines the snapback onto the baseline saltatory movement (Figure 3A, bottom left), in turn determining changes in distance traveled and period of saltatory movement in the presence of signal.

**Figure 3:**
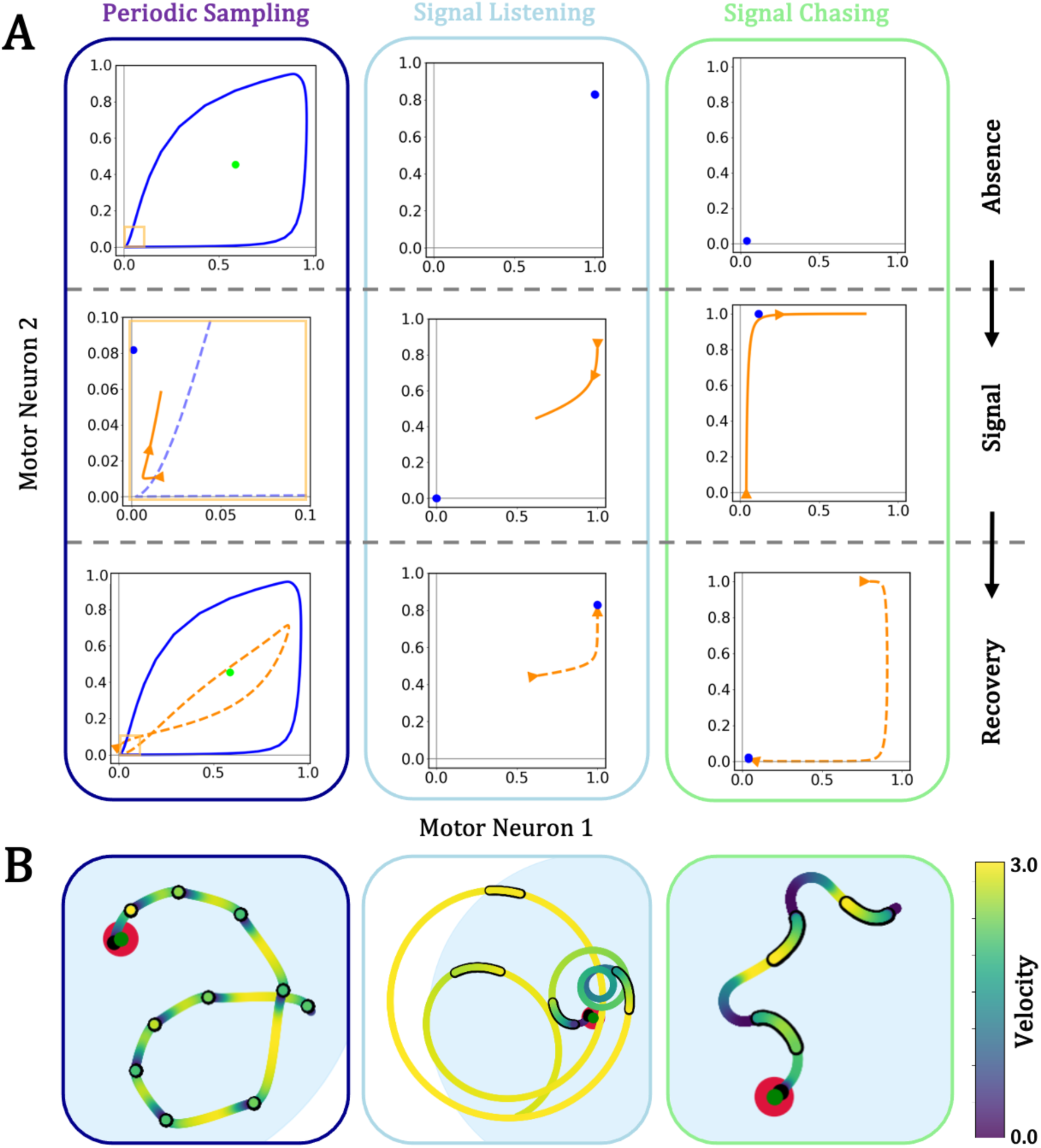
Three mechanisms that relate target signal and movement in evolved foragers. **(A)** Nervous system equilibria and dynamics projected to the motor neurons. In each panel, blue and green objects denote stable and saddle limit sets while orange traces show sample motor dynamics in the presence (solid) and absence (dashed) of target signal. (Left) Foragers that periodically sample converge on a cyclical motor pattern of fast and slow movement, yielding saltatory movement independent of signal. The onset of signal pulls nervous dynamics off of that baseline cycle, which can result in either an increase (signal chasing) or decrease (signal listening) in movement as the nervous dynamics reconverge to the intrinsic limit cycle. Note that here the presence of signal panel is zoomed in (yellow highlight). (Center) In signal listening, the forager has intrinsically high motor output, resulting in a cruise state (blue attractor). The onset of signal inhibits motor neurons, and motor dynamics approach the new attractor location at a complete stop. When the signal is removed, motor dynamics recover to the original attractor. (Right) In signal chasing, the opposite relationship occurs. The forager has intrinsically low motor output. The onset of signal excites the motor neurons, and signal loss returns nervous dynamics back to the slower state. **(B)** Sample behavior demonstrating each mechanism. Black outlined and non-outlined traces denote position with and without signal respectively. The green point indicates the target, the blue circle the detection radius, and the red circle the interaction radius. Traces are colored by forager velocity.

#### Signal Listening

Signal listening, the depression of motor activity in the presence of a target signal, was identified in both periodic and non-periodic foragers. In non-periodic foragers, signal listening generates saltatory motion by depressing movement relative to a faster cruising behavior in the absence of signal. Cruise movement is driven by a fixed nervous equilibrium with high motor output (Figure 3A, top center). The introduction of signal drops that nervous equilibrium to have little to no motor output (Figure 3A, middle center). In turn, the forager slows significantly until the loss of signal reintroduces the original nervous equilibrium and forager speeds back up (Figure 3A, bottom center). Forms of signal listening were identified in *UI, RD,* and *UD* foragers depending on the distance and relative heading to the target. This mechanism should not be expected to yield any regular period or duration of movement phases as change in movement is entirely driven by the signal pattern. In the case of periodic sampling, signal listening will likewise cause deviation from a regular period rather than enforcing one.

#### Signal Chasing

Signal chasing, the excitement of motor activity in the presence of a target signal, was also identified in both periodic and non-periodic foragers. Signal chasing is the antithesis of signal listening; signal chasers ambush the signal when it is available and otherwise remain relatively sedentary. Like signal listening, signal chasing occurred in both periodic and non-periodic foragers. That sedentary stage is based on a nervous equilibrium with low to negligible motor output (Figure 3A, top right). The introduction of signal excites activity the motor neurons (Figure 3A, middle right). Upon loss of the signal, the nervous dynamics return to the original rest state (Figure 3A, bottom right). Forms of signal chasing were identified in *UI, RD,* and *UD* foragers depending on the distance and relative heading to the target. Like signal listening, signal chasing should not be expected to yield any regular period or duration of movement phases as change in movement is entirely driven by the signal pattern. In the case of periodic sampling, signal chasing, like signal listening, will cause deviation from a regular period rather than enforcing one.

#### Inter and Intra-Individual Variation in Pursuit Strategy

Periodic sampling, signal chasing, and signal listening varied both within and between individual evolved foragers. Periodic sampling was determined categorically by the presence of an intrinsic limit cycle. Periodic sampling appeared in all but one *RD* forager (98.6%) and all *UD* foragers (100%). Signal listening and signal chasing were observed in *RD*, *UD*, and *UI* foragers. Both signal listening and chasing were quantified through a perturbation analysis with slightly different measures for periodic and non-periodic foragers. Signal listening and signal chasing varied in magnitude and often were both exhibited in individual foragers depending on distance from and heading relative to the target. Here, perturbation results are summarized for non-periodic and periodic foragers (see also Table 1).

#### Non-Periodic Foragers

Non-periodic foragers only evolved with unreliable targets (*U*). As such, for systematic combinations of target distance and relative heading, non-periodic foragers were exposed to a signal of the same length they experienced in evolution (*s_P_ * T/ s_I_*). Behavior and neural data before, during, and after the signal were recorded. To determine the magnitude of signal chasing and listening, each perturbation was compared to null behavior in which no signal was ever introduced. Measured responses to the signal were bounded by the onset of the signal and the earliest timestep where the forager returned to its baseline behavior, determined by when the motor neurons had returned to the values they held before signal onset (within 0.01 for both *x_1_* and *x_2_*). Duration between signal onset and return to baseline behavior and the resulting difference in distance compared to the null over the same duration were both collected (Figure 4A, left). Velocity change during the response could then be calculated, with acceleration indicating signal chasing and deceleration indicating signal listening.

**Figure 4:**
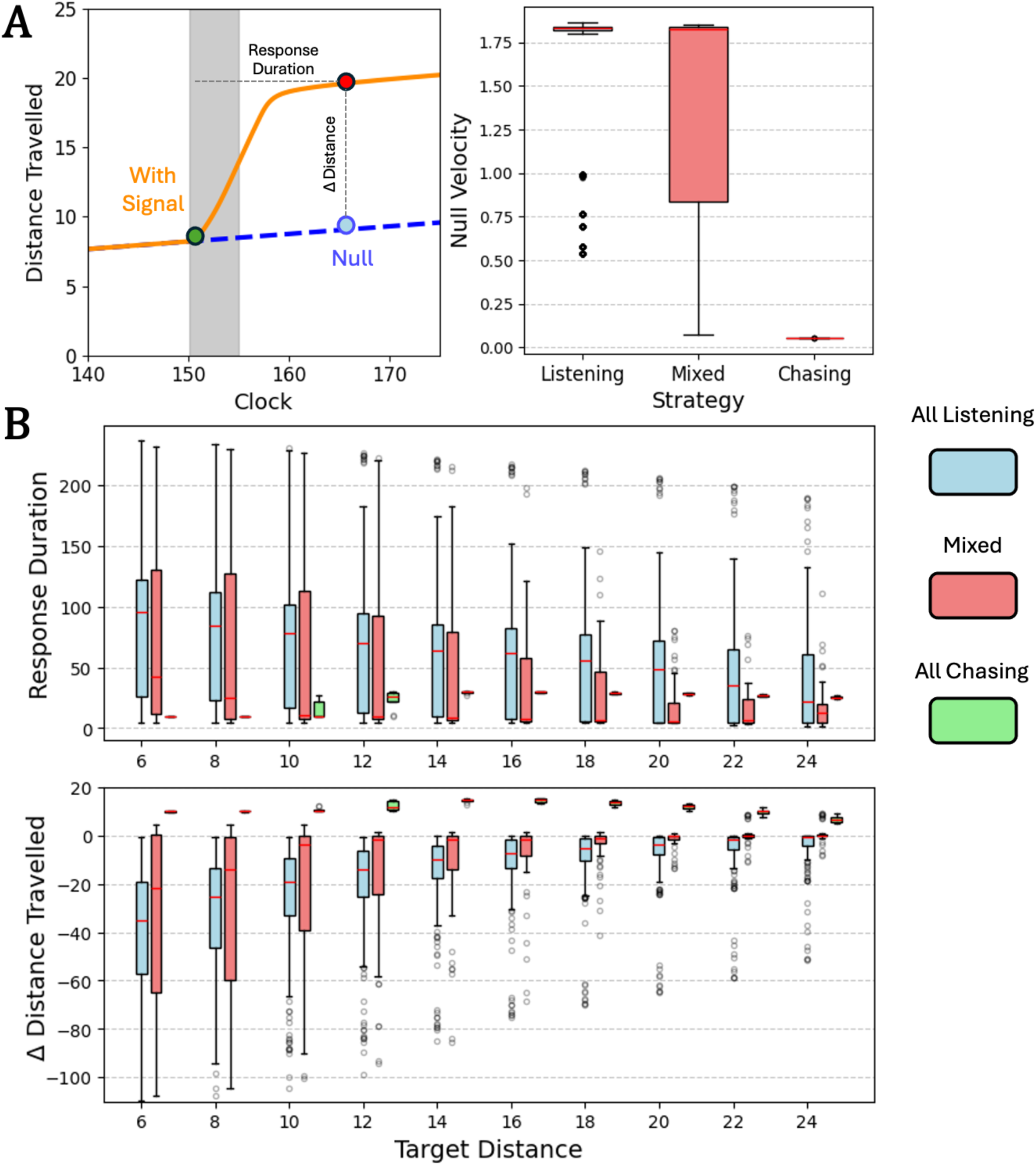
Non-periodic forager response to signal as a function of distance from their target. Blue boxes indicate foragers that only employed signal listening during pursuit, green those that employed only signal chasing, and pink those that relied on some mix of the two. Boxes indicate interquartile ranges, while red lines indicate means. **(A)** (Left) Non-periodic strategies were measured via the duration of response or the time between signal onset (green point, with signal presence denoted by gray bar) and return to pre-signal behavior (red point) and the resulting difference in distance travelled compared to the same time interval with no signal ever introduced (blue point). (Right) Without signal, foragers that only employed signal listening had significant null velocities in the absence of signal (mean at ∼41% of the maximum), those that only employed signal chasing had negligible velocity without signal, and those with mixed strategies demonstrated a spread between those extremes **(B)** Duration of forager response and change in distance travelled across target distance. Data is displayed for all different initial target headings for each agent.

Non-periodic foragers were made up of the entire *UI* population and one *RD* forager. Most *UI* foragers (67.3%) relied only on signal listening during pursuit. This majority exhibited a cruising velocity around half the maximum (δ/ *f*) in the absence of signal (Figure 4A, right). Duration of the movement response during signal listening varied depending on distance from the target. The magnitude of signal listening responses increased the closer *UI* foragers were to their targets, with longer response duration and shorter distance travelled (Figure 4B, blue boxes). In effect, this allows foragers to remain closer to their target awaiting the onset of the next signal. Conversely, the one non-periodic *RD* forager (1.4% of *RD*) and one *UI* forager (2.2% of *UI*) only exhibited signal chasing during pursuit. Both these foragers exhibited negligible velocity in the absence of signal (Figure 4A, right). Upon receiving signal, these foragers made modest jumps for short durations and were especially conservative both close to the target and close to the edge of the detection radius which here is set to 29 (Figure 4B, green boxes). Several other *UI* foragers adopted mixed strategies (30.5%) that used some combination of signal listening and signal chasing depending on the distance to and orientation from the target. These mixed foragers exhibited null velocities in the absence of signal that covered the range between that of the signal listeners and signal chasers (Figure 4A, right). Mixed strategies generally skewed towards signal listening but were more conservative and demonstrated lower mean durations across the board (Figure 4B, pink boxes).

#### Periodic Foragers

The perturbation analysis was different for periodic foragers. Periodic foragers were exposed to signal the first time they sampled for signal (*v* ≤ *ϕ*) after the transient period. Their subsequent behavior and neural data were recorded as in the non-periodic case. Three measurements were taken for periodic foragers: how long the forager sampled (*v* ≤ *ϕ*; *Δ* sampling time), the movement period from signal onset to the next sample (*Δ* period), and the resulting distance travelled (*Δ* distance; Figure 5A, left). Once again, each perturbation recording was compared to a null recording in which no signal was introduced. In this case, signal chasing would be identified by acceleration leading to shorter sample times (since sampling stops once *v* > *ϕ*) and movement period durations, with resulting decreases in distance travelled. Conversely, signal listening occurs with deceleration and longer sample times and movement period durations, with resulting increases in distance travelled.

**Figure 5:**
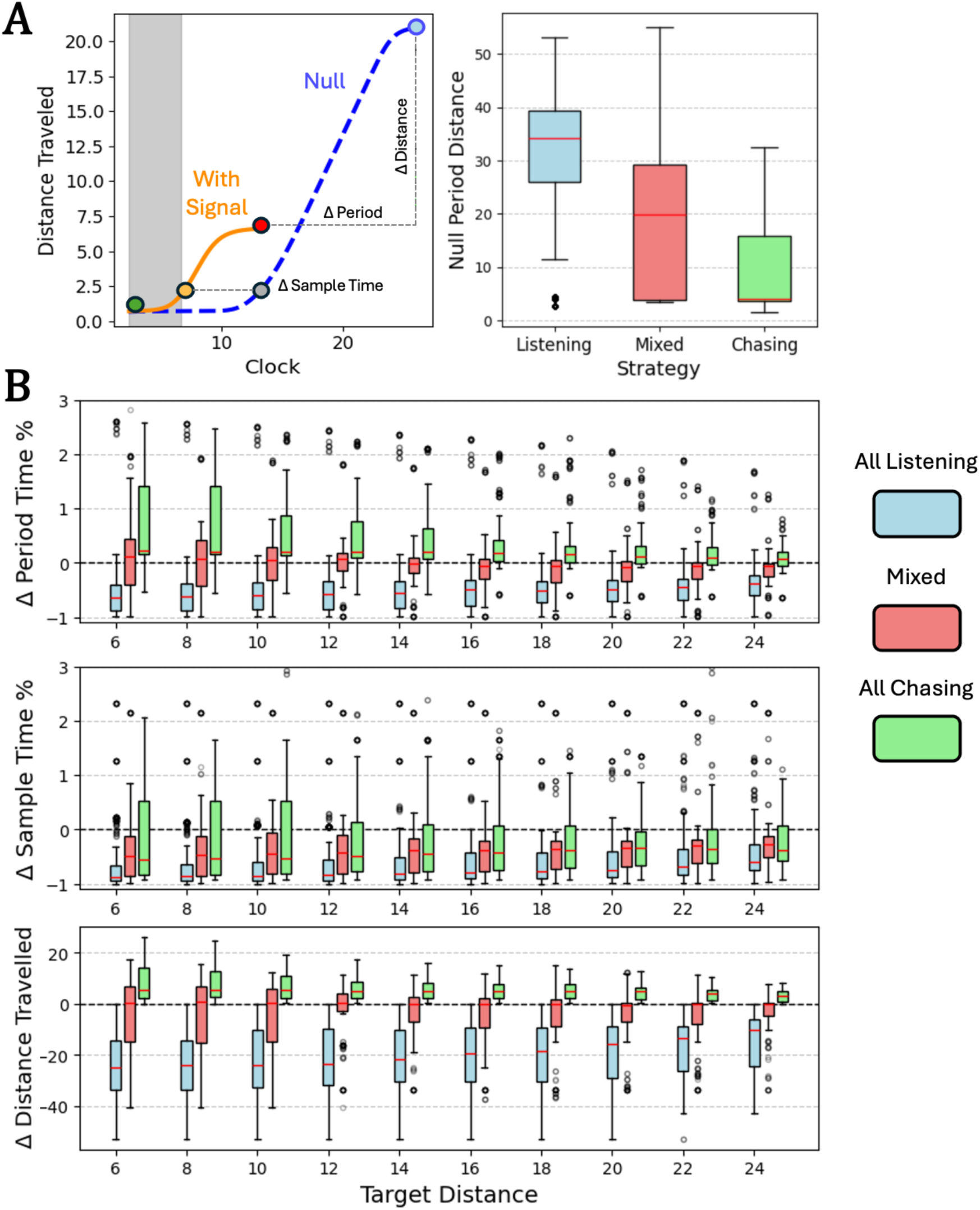
Non-periodic forager response to signal as a function of distance from their target. Blue boxes indicate foragers that only employed signal listening during pursuit, green those that employed only signal chasing, and pink those that relied on some mix of the two. Boxes indicate interquartile ranges, while red lines indicate means. **(A)** (Left) Periodic strategies were measured via the proportional change of the period (green point to red point compared to green point to blue point, with signal presence denoted by gray bar), the proportional change of the sample time (*v ≤ ϕ*; green point to orange point compared to green point to gray point), and the resulting difference in distance travelled from the null (y-difference of red and blue points). (Right) Without signal, foragers that only employed signal listening had longer movement distances each period than those that only employed signal chasing. **(B)** Period change, sample time change, and change in distance travelled across target distance. Data is displayed for all different initial target headings for each agent.

Periodic foragers were made up of the entire *UD* population and all but one *RD* forager. Unlike non-periodic foragers, periodic foragers that only relied on signal listening were a minority (11.6% of *RD*, 10.8% of *UD*). This minority exhibited sizable movement periods in the absence of signal (Figure 5A, right). Once again, duration of movement and sampling time during signal listening depended on distance from the target, although responses were generally lower in magnitude than those of non-periodic foragers. Both period duration and sampling time declined the closer signal listeners were to their targets, leading to higher frequency sampling at closer proximity and shorter distances travelled between those samples (Figure 5B, blue boxes). Like their non-periodic counterparts, this gives periodic signal listeners a better chance of remaining close to their target when they receive the next signal. Periodic foragers exhibited solely signal chasing strategies far more than their non-periodic counterparts (56.5% of *RD*, 45.9% of *UD*). Signal chasing periodic foragers generally had short period and thus high frequency sampling strategies in the absence of signal (Figure 5A, right). Upon receiving signal, movement periods were generally longer for these foragers, although these differences were very slim for the most part (Figure 5B, green boxes). As such, the baseline high frequency behavior is likely of more consequence to this strategy than adjustments made upon signal detection. Finally, like the non-periodic case, several periodic foragers used mixed strategies (31.9% of *RD*, 43.5% of *UD*) depending on target distance and orientation that interpolated between the signal listening and signal chasing extremes. These mixed foragers exhibited null periods that covered the range between that of the periodic signal listeners and signal chasers (Figure 5A, right). On average, mixed periodic strategies did not skew strongly toward either signal listening or signal chasing (Figure 5B, pink boxes). The only reliable relationship in mixed strategies was a shorter sampling time in the presence of signal, although this was mostly true for pure signal listeners and chasers as well.

## Discussion

Our simulations show that foragers’ detection of signals can generate saltatory patterns of movement during pursuit that shift according to the distance from a signaling target. Foragers receiving reliable signals and having perception that worked independent of their own movement (*RI*) employed cruise-like pursuit strategies. However, when forager perception was dependent on movement, they adopted saltatory movement strategies both with reliable (*RD*) and unreliable (*UD*) signals. Strategies evolved in which signals altered the period and subsequent distance of saltatory movement in both these cases. Generally, periodic foragers, which moved in a saltatory manner even in the absence of signal, were less sensitive to target signals than non-periodic foragers, balancing the need to move with the need to detect the target. For non-periodic foragers, the patterns of signal alone generated saltatory pursuit movements by either exciting (signal chasing) or inhibiting (signal listening) movement. Signal chasing generally results in more restricted movement in the absence of signals, especially close to the edge of the detection radius. Conversely, signal listening leads to more conservative movement in the presence of signals and closer to the target. In both cases, the exact character of the resulting saltatory movement depends on the pattern of signal a forager receives from the target.

Observing regular periodic saltatory motion during search may suggest that some sensory limitation is generating the saltatory movement, necessitating stops to enable detection of targets. Trade-offs between perception and movement during pursuit are common in the animal kingdom, especially in cases of non-visual detection of signals. For example, periodic sampling is the norm for animals that rely on tactile perception (Bassett et al., 2007). Benthic-dwelling fish orient to and strike tactile stimuli during pursuit through a series of punctuated movements and samples along the benthic substrate (Hoekstra & Janssen, 1985; Janssen, 1990; Janssen & Corcoran, 1998; Kornis et al. 2012). The bursting approach of these fish to their targets provides the fish information about their target in a ‘stepwise’ manner during pursuit (Coombs & Conley, 1997; Bassett et al., 2007). Wading birds similarly take advantage of tactile stimuli using “remote touch” to detect prey at night, in high turbidity, or when prey is buried (Piersma et al. 1998; Cunningham et al. 2009; Cunningham et al. 2010), yielding periodic sampling patterns whose period depends on previously detected prey signals (e.g., Dias et al. 2009). Olfactory foraging and chemotaxis more generally are another case where periodic sampling is commonly demonstrated (Berg, 1975; Lockery, 2011; Gomez-Marin et al. 2014; Svensson et al. 2014; Baker et al. 2018). In all these cases, it would be interesting to test our findings here of how periodic pursuit movements increase in frequency and decline in distance with increased proximity to a target.

Saltatory patterns of movement may also emerge in otherwise cruising or stationary animals post-encounter due to intermittent signaling by unreliable targets. The most common mechanism of saltatory movement in model foragers was signal listening, the depression of movement in the presence of a target signal. Localization of movement during search is well documented, commonly referred to as ‘area-restricted-search’ (ARS, Tinbergen et al. 1967; Kölzsch et al. 2015). ARS describes the ability to plastically alter search effort depending on encounters with targets or some indication of habitat quality (Dorfman et al. 2022). In most cases, the initial detection of a target signal breaks an otherwise continuous or cruise search strategy and yields a slowing of movement at the onset of ARS. That said, ARS does not speak to pursuit. A similar pattern of movement depression from signal can occur during pursuit and can be an effective movement strategy after an encounter has occurred. We also found that movement depression during pursuit often strengthened in proximity to a target for both periodic and non-periodic foragers.

The inverse strategy, signal chasing, was also a viable pursuit strategy for the simulated foragers. It could be argued that the ambush foraging mode captures this behavior, but ambush strategies are generally described as one-shot movements where a single burst of movement entails both pursuit and capture. Many animals certainly rely on this ambush foraging mode, but here signal chasers demonstrate that saltatory movement may result from repeated “ambushes” of signals throughout pursuit. Signal chasing may occur when the target of search is sessile or moves much more slowly than its predator, but its signals are not easy to triangulate, such as the pursuit of eggs (e.g., Gervais et al., 2021; Kostoglou et al., 2021) or that for buried invertebrates by birds (e.g., robins and worms – Montgomerie & Weatherhead, 1997). The foraging animal may thus only move when signal is available, for fear of drifting away from the target in the signal’s absence. Our simulated foragers showed that this effect may be especially strong close to the target or conversely at the edge of the forager’s perceptual range. Signal chasing may also be more likely when the target is rare and losing signal from an encountered target is more detrimental.

More generally, pursuit is often a more complicated problem than a simple race. Here, evolved foragers often relied on multiple movement strategies depending on their spatial relationship to their targets and the motion-dependence of their perception. There are other pursuit circumstances that call for further explanation. Pursuit of a moving target is a rich area that has been explored at length (Isaacs 1999; Nahin, 2012; Weintraub et al. 2020) and recent work has further demonstrated perception as a major determinant of pursuit character and success (Martin et al. 2022). However, like the case of a stationary target, pursuit of a mobile target with unreliable signal is not as well characterized. Furthermore, target signal patterns are likely to evolve specifically to confound pursuers. Our simple sessile target task could also be used to explore how foraging movement strategies may develop in cases where the signal from the target is not random but rather controlled by the target to best confound the current strategy of the forager. Co-evolution of mobile pursuit has been explored before (Nolfi & Floreano, 1998), but modeling co-determined signal and movement strategies between forager and target is an important research direction as they are more likely to reflect real predator-prey behavioral strategies.

We believe the mechanisms of variation in pursuit movement explored here can help demystify naturalistic animal movement patterns as data about those patterns continues to proliferate. Models of the kind we present here afford many possibilities for exploring the problem space of signal perception during foraging. These models also could provide further insight into the composition of movement at different stages of the foraging cycle. More modeling efforts of this kind are thus called for to help generate and supplement a more coherent theory of animal movement.

## Appendix

### Section 1: Model Parameters

| Parameter | Notation | Value |
| --- | --- | --- |
| **Task Parameters** |  |  |
| Duration | $T$ | 600 |
| Transient Duration | $T_t$ | 100 |
| Step size | - | 0.05 |
| Friction coefficient | $f$ | 0.9 |
| Initial target distance | $r_0$ | 23 – 27 |
| Interaction threshold | $\lambda$ | 2 |
| Unreliable signal availability | $s_P$ | 0.3 |
| Unreliable signal appearances | $s_I$ | 20 |
| **Forager Parameters** |  |  |
| Detection radius | $d$ | 30 |
| Sensor offset | $o$ | 2 |
| Maximum thrust | $\delta$ | 2 |
| Maximum torque | $\varepsilon$ | $\pi/6$ |
| Maximum sensor value | $\gamma$ | 3 |
| Movement constraint | $\phi$ | 0.01 |
| **CTRNN Parameters** |  |  |
| $\mathbf{x}$ range | - | 0 – 1 |
| $\boldsymbol{\tau}$ range | - | -16 – 16 |
| $\boldsymbol{\theta}$ range | - | -16 – 16 |
| $\mathbf{a}_{ji}$ range | - | -16 – 16 |
| $\mathbf{b}_{ji}$ range | - | -20 – 20 |
| **Evolutionary Algorithm Parameters** |  |  |
| Population size | - | 400 |
| Generations | - | 1000 |
| Mutation variance | - | 0.12 |
| Elitism | - | 0.02 |
| Crossover | - | 0.01 |

### Section 2: Robustness Analysis

Robustness analysis was performed on each of the successful agents (86 *RI*, 46 *UI*, 69 *RD*, and 37 *UD*) to determine performance within the trained range of initial positions (3 – 7 units from the edge of detection radius *d*) and performance extrapolated at other positions within *d*. Robustness analysis of this kind is often useful in evaluating the generality of evolved solutions when using evolutionary algorithms. Behaviors that do not extrapolate outside the initial conditions used during evolution may be overfit to those conditions and not represent a general solution. As such, we evaluated each agent’s fitness (eq. 5, main text) at each of 5400 initial locations within detection radius *d*. 900 were within the evolved range of initial positions while 4500 extrapolated outside of that range. Fitness at each location was averaged for five simulations starting with different initial headings to the target (intervals of 2π/5).

The performance of the best performing forager in each condition over the course of each evolutionary run shows how performance on the pursuit task changed during evolution (Figure A1, left). Robustness analyses for the best forager in each of those populations (Figure A1, A, *RI*; B, *UI*; C, *RD*; D, *UD*) is exemplified by coloring each initial position by the trial fitness at that position. For the most part, evolved pursuit strategies generalized well outside of the evolved initial conditions.

**Figure A1:**
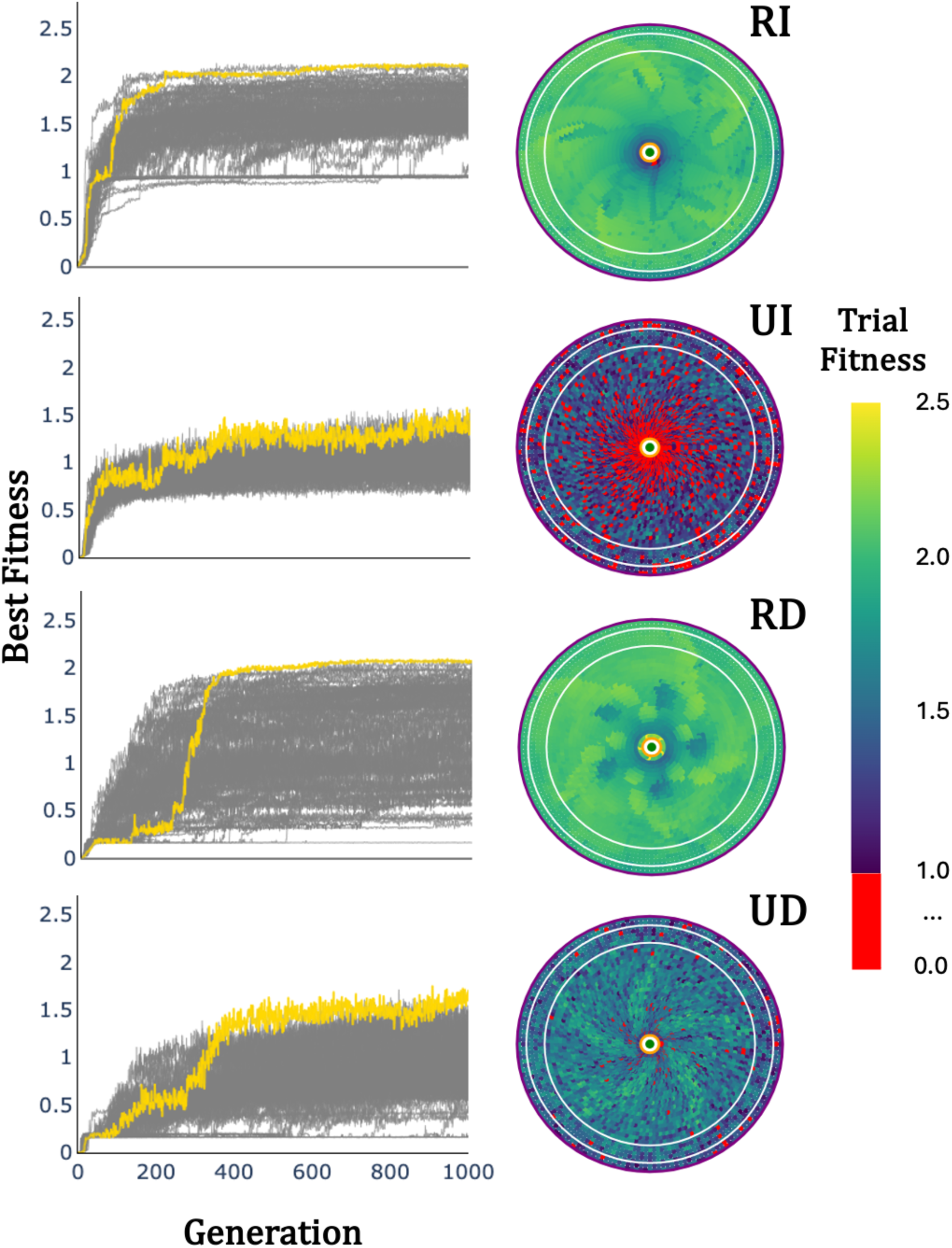
Evolution and performance robustness of highest fitness foragers in 100 evolutionary runs of each condition. Left panels show the evolutionary progress of the best individual in each evolution, with the best overall individual highlighted in gold. The rights panels show the average fitness across five initial headings of that best overall individual initialized from various points within range of the target. Foragers were evaluated during evolution only from initial conditions between the two white lines, all other points show the success (or lack thereof) of a forager starting in that position (averaged over five initial orientations). The detection radius of the forager is denoted in purple while the interaction radius is shown in orange. All successful trials (F >= 1) are colored according to fitness while unsuccessful trials are colored red. **(RI)** reliable targets, movement-independent perception, **(UI)** unreliable targets, movement-independent perception, **(RD)** reliable targets, movement-dependent perception, and **(UD)** unreliable targets, movement-dependent perception.

## Notes

### Competing Interest Statement

The authors have declared no competing interest.

